# Response to SMAC mimetics in patient-derived models of triple-negative breast cancer relies on RIPK1 activation

**DOI:** 10.64898/2026.10.26.761582

**Authors:** Elisabeth Brown, Paige Jensen, Chieh-Hsiang Yang, Emilio Cortes Sanchez, Zhengtao Chu, Huiyi Yang, Yi Qiao, Martin Golkowski, Apurva Srivastava, Bryan E. Welm, Jaime Fornetti, Alana L. Welm

## Abstract

Triple-negative breast cancer (TNBC) is a highly metastatic and aggressive breast cancer subtype with limited targeted treatment options. Second mitochondrial activator of caspases (SMAC) mimetics represent a promising therapeutic class for TNBC patients, although they have yet to be approved clinically. Screening a repository of 69 breast cancer patient-derived xenograft organoid (PDxO) lines revealed that approximately 39% were sensitive to the SMAC mimetic birinapant. To define the basis of resistance in lines that did not respond, we compared birinapant-sensitive (S-) and *de novo* resistant (R-) TNBC PDxOs. Birinapant caused degradation of inhibitor of apoptosis proteins (IAPs) in both groups but failed to induce cell death in R-PDxOs. We determined that R-PDxOs were susceptible to certain types of apoptosis but were specifically resistant to TNFR-mediated cell death, an integral pathway involved in SMAC mimetic response. Moreover, RIPK1 activation via autophosphorylation, a critical event in TNFR-mediated apoptosis, was absent in both R-PDxOs and PDxOs selected for acquired birinapant resistance. Pharmacologic inhibition or degradation of RIPK1 protected S-PDxOs from birinapant-induced cell death, demonstrating that RIPK1 activation is required for response in TNBC. To investigate how to restore RIPK1 activation in R-PDxOs, we discovered that HOIP, an E3 ubiquitin ligase that inhibits RIPK1 autophosphorylation, was more abundant in R-PDxOs compared to S-PDxOs. Inhibiting HOIP activity restored RIPK1 autophosphorylation and sensitized a subset of R-PDxOs to birinapant. Together, these findings identify impaired RIPK1 autophosphorylation as a mechanism of birinapant resistance in TNBC and suggest that RIPK1 activation state may distinguish SMAC mimetic-sensitive and -resistant tumors. Targeting HOIP may provide a strategy to overcome resistance in selected tumors.

## Introduction

In the United states, breast cancer is the second most frequently diagnosed cancer and the second leading cause of cancer-related deaths in women^1^. The triple-negative breast cancer (TNBC) subtype, which exhibits low or absent expression of the estrogen receptor (ER), progesterone receptor (PR), and human epidermal growth factor receptor 2 (HER2), has the lowest 5-year relative survival rate and is the most likely to recur early as a distant metastasis^2,3^. Upon diagnosis, standard treatment for TNBC includes neoadjuvant chemotherapy and immune checkpoint inhibitors, surgery and radiation. Additional chemotherapy or immunotherapy is given as adjuvant therapy for high-risk cases^3,4^. For recurrent metastatic TNBC, treatment options include antibody-drug conjugates, additional chemotherapy, and certain targeted therapies approved depending on presence of mutations^3,4^. However, resistance to all available therapy is common and the prognosis for metastatic TNBC is poor^5,6^. Thus, compared to other breast cancer subtypes, TNBC has more limited treatment options^7^, highlighting the need for new therapeutic strategies to improve patient outcomes.

One potential new class of targeted therapeutics for TNBC is secondary mitochondrial activator of caspases (SMAC) mimetics. SMAC mimetics imitate cellular SMAC function by targeting inhibitor of apoptosis proteins (IAPs) for degradation, initiating cell death signaling^8–10^. In the intrinsic apoptosis pathway, SMAC mimetics degrade x-linked IAP (XIAP) and induce cleavage of caspase-9, -3, and -7, while in the tumor necrosis factor receptor (TNFR) extrinsic apoptosis pathway, they degrade cellular IAPs 1 and 2 (cIAP1/2)^8–10^. cIAP1/2 are E3 ubiquitin ligases that can add K63-linked ubiquitin to target proteins to promote their scaffolding function, which determines the signaling program executed downstream of the TNFR – specifically, NF-κB signaling, apoptosis, or necroptosis^8–13^. When cIAP1/2 are degraded, proteins involved in cell death signaling, namely RIPK1 and TRADD, untether from the TNFR and couple with proteins in the cytoplasm to form cell death complex IIb (RIPK1, TRADD, FADD, caspase-8) or the necrosome (RIPK1, RIPK3, MLKL), depending on caspase-8 presence^8–12,14^. Complex IIb formation results in apoptosis via cleavage of caspases-8, -3, and -7, whereas the necrosome induces necroptosis independently of caspases^8–11,14,15^. SMAC mimetics have been previously tested in breast cancer patient-derived xenografts (PDX), PDX-derived spheroids, and standard breast cancer cell lines, revealing that some tumors are sensitive to SMAC mimetics but others are not, despite successful drug delivery and intended on-target effects^14,16^.

Although the mechanism of action of SMAC mimetics has been characterized extensively and some preclinical models show evidence of response, the clinical efficacy of these drugs has been disappointing. According to ClinicalTrials.gov, over 40 clinical trials have been initiated to evaluate SMAC mimetics across multiple cancer types, including three trials that included TNBC patients. Notably, in one phase II trial (NCT01617668), patients with localized TNBC were randomized to receive either the chemotherapeutic paclitaxel or paclitaxel in combination with the SMAC mimetic LCL161. Despite unexpected toxicity of paclitaxel combined with LCL161, and no observed difference between treatment groups, a prospectively planned analysis revealed that patients within the combination treatment group whose tumors had higher expression of a pre-determined TNF-α-based gene signature showed a 21% higher rate of pathological complete response (pCR) compared to those with lower expression^17^. These clinical results suggest that the TNBC patient population comprises distinct subsets of SMAC mimetic responders and non-responders that require additional investigation to identify. Although the NCT01617668 trial demonstrated a potential gene signature associated with LCL161 response in TNBC, the molecular determinants of SMAC-mimetic resistance remain incompletely defined. Given that multiple phase II SMAC mimetic trials failed to demonstrate efficacy over standard of care, it is likely that these patient populations include tumors with intrinsic resistance to SMAC mimetics. Understanding the underlying biological mechanism(s) determining resistance may help explain the limited clinical efficacy and inform future strategies for patient selection and clinical trial design.

Since SMAC mimetics have been reported to successfully degrade IAPs as intended in patient tumors and preclinical models that do not respond to treatment^9,14,17^, we hypothesized that TNBCs resistant to SMAC mimetics fail to execute the cell death signaling cascade downstream of IAP degradation. Here, we utilized TNBC patient-derived models displaying SMAC mimetic sensitivity, *de novo* resistance, and acquired resistance to investigate the mechanism of resistance to birinapant, a SMAC mimetic whose mechanism of action has been previously characterized in TNBC^9,14–16^. We discovered that birinapant-resistant models are defective for activation of RIPK1 by autophosphorylation during birinapant treatment, and identified HOIP, the catalytic subunit of LUBAC, as a regulator of this response in the three resistant TNBC models tested. Overall, these findings establish dysregulated control of RIPK1 activity as a critical determinant of birinapant response in TNBC.

## Results

### SMAC mimetic treatment elicits a dichotomy of responses in triple-negative breast cancer PDxOs

We previously demonstrated that birinapant treatment of 16 breast cancer patient-derived xenograft organoids (PDxOs) elicited a range of responses, which was validated in a selected cohort of matched patient-derived xenografts (PDXs)^18^. To identify how effective SMAC mimetics were across a larger cohort of breast cancer patient-derived models that represent all clinical subtypes of breast cancer, those from primary or metastatic sites, and tumors collected pre- or post-treatment, we treated 69 PDxO models with birinapant for six days and assessed cell viability. Some organoid lines appeared resistant since they continued to grow in the presence of drug, whereas birinapant was cytotoxic to other lines, indicating drug sensitivity (**Fig. 1a**). To categorize sensitive versus resistant PDxOs based on a clinically relevant metric, we mapped the in vivo responses of our previously published cohort of PDX models treated with birinapant^18^ onto their matched PDxO (marked with asterisks, **Fig. 1a**), using the criteria of PDX clinical response (mRECIST) developed by the NCI PDX Network to define sensitivity and resistance^19^. Sensitive PDXs demonstrated a complete response (CR), whereas resistant PDXs showed progressive disease (PD)^18^. Treatment of our large cohort of PDxOs with birinapant elicited a similar range of responses as we observed in the smaller PDxO cohort with 39% of models demonstrating sensitivity and the rest exhibiting *de novo* resistance (**Fig. 1a**). Most, but not all, sensitive models were of the TNBC subtype. Evaluation of responses to five different SMAC mimetics showed a similar response profile across an expanded cohort of 20 TNBC PDxOs, containing one additional line (**Fig. 1b**). Thus, multiple SMAC mimetics produced concordant response profiles across a large cohort of patient-derived breast cancer models, with a subset exhibiting sensitivity. We next sought to elucidate the biological basis of this observed divergent response and determine if there are ways to reverse resistance to enhance efficacy.

**Figure 1.**
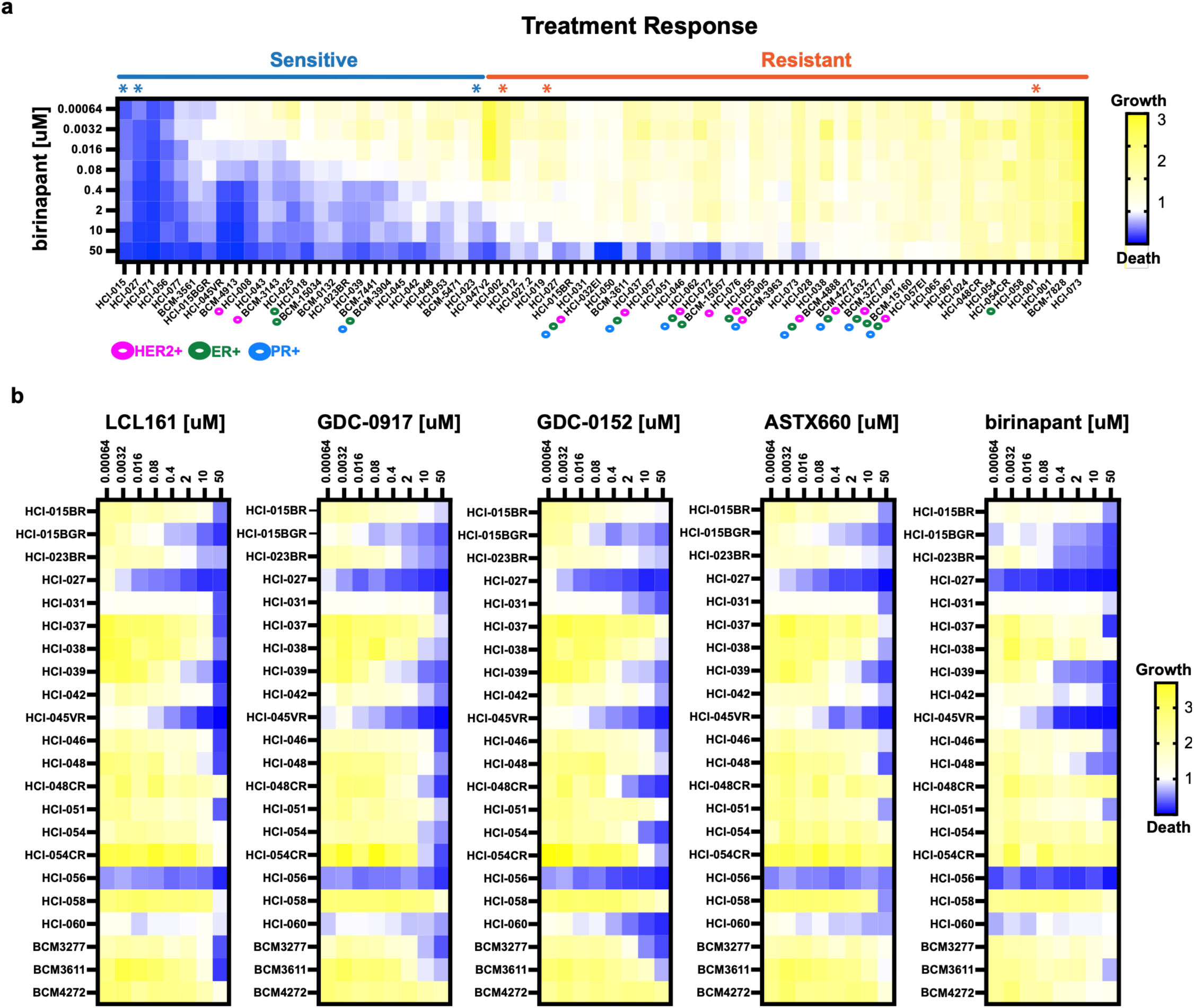
SMAC mimetic treatment elicits a dichotomy of responses in triple-negative breast cancer PDxOs. **(a)** Patient-derived xenograft organoid (PDxO) responses to 6 days of birinapant treatment at the indicated concentrations. Cell viability after treatment was measured via Cell Titer Glo (displayed as the average of 3 biological replicates with 4 technical replicates each). Asterisks (*) denote lines where response to birinapant was tested in vivo using the matched PDX model (see ref. 18). Expression of HER2, estrogen receptor (ER), and progesterone receptor (PR), as well as birinapant response status is also denoted. **(b)** Comparison of PDxO responses after treatment with 5 different SMAC mimetics for 6 days at the indicated concentrations, measured via Cell Titer Glo (average of 3 biological replicates with 4 technical replicates each). Luminescent values for each technical quadruplicate were normalized to their average day 0 control values and each set of normalized quadruplicates was averaged together to obtain one value per biological replicate. For all Cell Titer Glo heatmaps, blue indicates cytotoxicity, white indicates a cytostatic response, and yellow indicates growth compared to Day 0.

### Birinapant-resistant PDxOs are not impervious to other inducers of apoptosis

To verify that treatment effectively targeted IAPs in resistant lines, we treated birinapant-sensitive (S-) and birinapant-resistant (R-) TNBC PDxOs (two lines each) with birinapant for 12 hours and compared IAP protein levels between groups by Western blot. To ensure that the PDxO findings reflected an in vivo resistance phenotype, R-PDxOs previously shown to display birinapant resistance in their matched PDX models were selected for evaluation^18^. In all PDxOs tested, birinapant treatment caused reduction of cIAP1, cIAP2, and XIAP, as expected (**Fig. 2a**). Thus, resistance could not be explained by lack of on-target drug activity, aligning with prior clinical results^17^. To confirm that apoptosis via caspase cleavage was the mechanism of birinapant-induced cell death in sensitive lines, S-PDxOs were treated with birinapant in the presence of the pan-caspase inhibitor Z-VAD-FMK. As expected, caspase inhibition blocked birinapant-induced cell death in S-PDxOs (**Fig. 2b**). As a follow up, S- and R-PDXs were treated with one dose of birinapant in vivo, and tumors were collected at several timepoints to assess caspase-3 activation. We observed significantly increased cleaved caspase-3 in S-PDXs following treatment, whereas the R-PDX showed no significant induction of cleaved caspase-3 (**Fig. 2c**). Together, these results demonstrated that birinapant-resistant models failed to induce apoptosis despite degradation of IAPs, suggesting there could be defects in apoptotic signaling that promote resistance.

**Figure 2.**
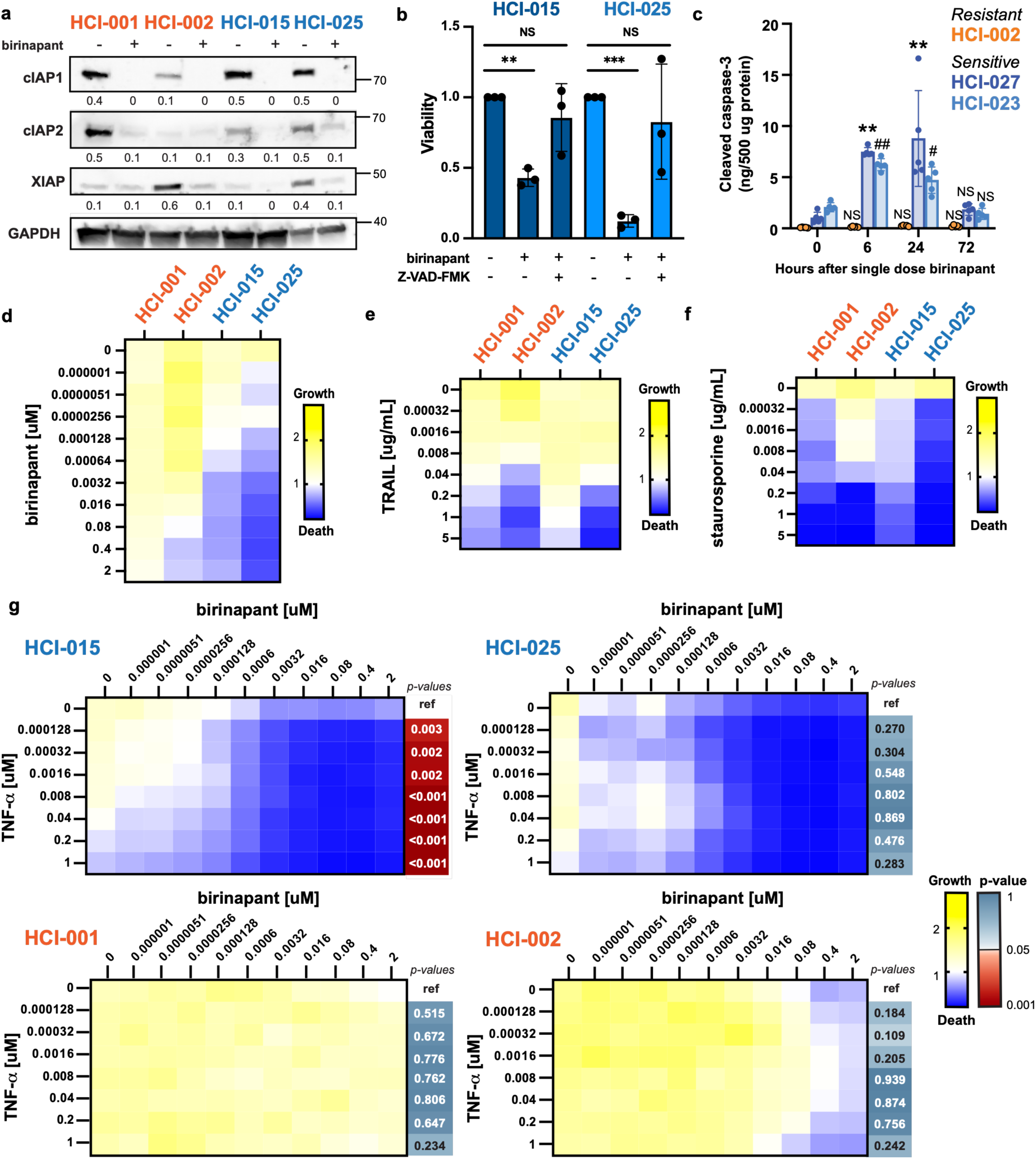
Birinapant-resistant TNBC models fail to induce apoptosis downstream of the TNFR. **(a)** SMAC mimetic-resistant (R-, orange) and -sensitive (S-, blue) TNBC PDxOs were treated with birinapant (10nM) for 12 hours in the presence of Z-VAD-FMK (25uM). IAPs were detected via Western blot. Abundance of each IAP was determined using densitometry and calculated as a ratio to GAPDH, indicated below each blot. **(b)** TNBC S-PDxOs were treated with birinapant (16nM) for 48 hours either in the presence of vehicle or Z-VAD-FMK (25uM). Cell viability after treatment was evaluated with Cell Titer Glo (3 biological replicates, with 4 averaged technical replicates each). Statistical significance was calculated (n=3 biological replicates per line) using a one-sample t-test, **p=0.004;***p=0.0009. NS, not significant. **(c)** TNBC R- and S-PDXs were treated with one dose of birinapant (n=5 per model, IP injection, 20mg kg^-1^) and tumors were collected at the indicated timepoints (0 is no treatment). The abundance of activated cleaved caspase 3 was measured in each tumor. Statistical significance was calculated using an ordinary two-way ANOVA with Dunnett’s multiple comparisons test, **p<0.0001 vs HCI-027 0 hour, ^#^p=0.0158 vs HCI-023 0 hour, ^##^p=0.0001 vs HCI-023 0 hour. Data are shown as mean +/- s.e.m. NS, not significant. **(d-f)** TNBC R- (orange) and S- (blue) PDxOs were treated with titrated birinapant, TRAIL, or staurosporine for 48 hours. Cell viability was evaluated via Cell Titer Glo (3 biological replicates with 4 technical replicates each). **(g)** TNBC R- (orange) and S- (blue) PDxOs were treated with combined titrations of birinapant and TNF-α for 48hr. Cell viability was evaluated with Cell Titer Glo (3 biological replicates with 4 technical replicates each). For each biological replicate, the area under the curve (AUC) for each combination treatment row was compared with the corresponding AUC for birinapant alone (row 0; ref = reference) to determine the effects of TNF-α on birinapant response. Statistical significance between AUCs was assessed using a two-sided paired Student’s t*-*test (n = 3 biological replicates), and p-values are color-coded by significance. For all heatmaps, darker blue indicates cytotoxicity, white indicates a cytostatic response, yellow indicates growth, and red p-values indicate statistical significance. Detailed statistics for single-agent heat map data and calculated AUC values for each biological replicate are shown in **Supp. Fig. 1**.

To determine the extent of apoptosis resistance in R-PDxOs, we stimulated R- and S-PDxOs with birinapant, staurosporine, or TRAIL to induce different apoptotic signaling pathways. Staurosporine has been shown to induce intrinsic apoptosis^20,21^, whereas TRAIL activates death receptor 5 (DR5) to induce extrinsic apoptosis independently of IAP degradation^22,23^. Evaluation of cell viability after treatment relative to the untreated control revealed that responsiveness to staurosporine or TRAIL did not correlate with birinapant resistance, and R-PDxOs were able to execute cell death using pathways that do not depend on the TNFR (**Fig. 2d-f**, statistics shown in **Supp. Fig. 1a-c**). This was consistent with prior literature demonstrating that the TNFR plays a critical role in response to SMAC mimetics, including birinapant^9,24^.

To dissect any differences in TNFR-mediated cell death between R- and S-PDxO lines, we titrated R- and S-PDxOs with birinapant, TNF-α, or the combination treatment to further stimulate TNFR-mediated apoptosis^14^. Analysis of cell viability and comparison of calculated area under the curve (AUC) values of birinapant response across TNF-α concentrations demonstrated that R-PDxOs were resistant to TNFR-mediated cell death, whereas apoptosis was enhanced or sustained in S-PDxOs (**Fig. 2g, Supp. Fig. 1d)**. Importantly, R-PDxOs displayed no significant differences in levels of proteins involved in TNFR signaling (cIAP1, cIAP2, TRAF2, RIPK1, TNFR1, TRADD, and FADD; **Supp. Fig. 2a**), and they retained their ability to induce NF-κB activation following TNF-α/TNFR stimulation (**Supp. Fig. 2b**). These data indicated that the TNFR was functional in R-PDxO lines; however, the observed resistance to cell death in response to TNFR-dependent apoptotic stimuli pointed to a defect downstream of TNFR that conveyed birinapant resistance in R-PDxOs.

### PDxOs resistant to birinapant are defective in RIPK1 autophosphorylation

RIPK1 is a key component of TNFR-mediated signaling^9,12^, and activation of RIPK1 by autophosphorylation at S161 and S166 is required for TNFR-mediated apoptosis^12,14^. To assess RIPK1 activation following birinapant treatment, we treated S- and R-PDxOs with birinapant and assessed RIPK1 phosphorylation at S166 as a marker for RIPK1 activation^25^, by Western blot. The result was striking: S-PDxOs induced RIPK1 phosphorylation by 6 hours after birinapant treatment, with accelerated peak induction in the presence of TNF-α (**Fig. 3a, top**). In contrast, R-PDxOs failed to phosphorylate RIPK1 at S166, even in the presence of exogenous TNF-α to increase TNFR signaling^14^ (**Fig. 3a, bottom**). We also observed concomitant reduction of total RIPK1 in S-PDxOs following birinapant treatment (**Fig. 3a, top**), which is indicative of RIPK1 cleavage by caspase 8 (**Supp. Fig. 2a**) after initiation of apoptosis, as previously described^26^. RIPK1 mRNA levels remained unchanged during birinapant treatment (**Supp. Fig. 2b**), consistent with post-translational regulation of RIPK1 protein.

**Figure 3.**
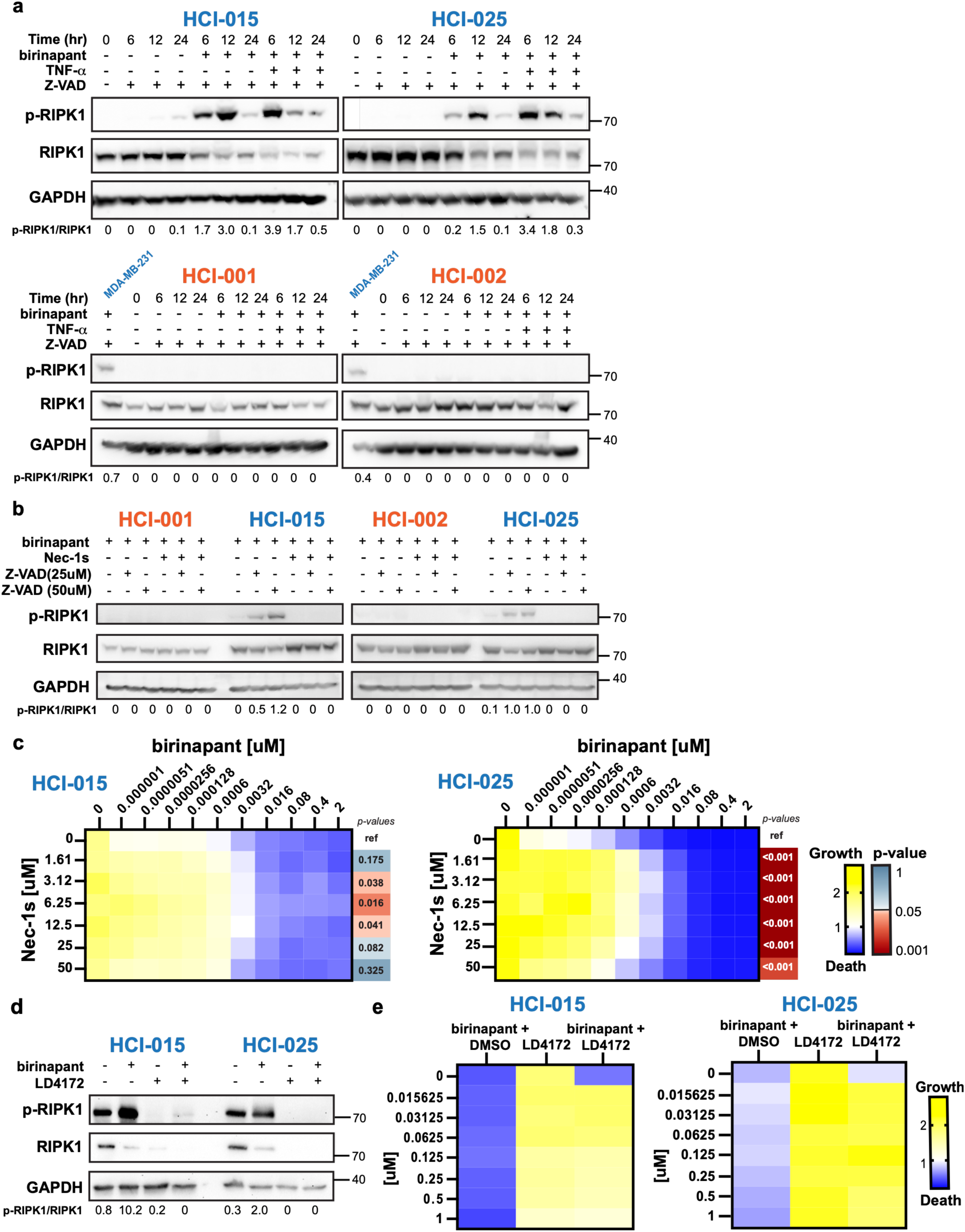
Birinapant-resistant TNBC PDxOs fail to activate RIPK1. **(a**) TNBC R- (orange) and S- (blue) PDxOs were treated with birinapant (10nM) +/- TNF-α (20ng/mL) in the presence of the caspase inhibitor Z-VAD-FMK (Z-VAD; 25uM) for the indicated time in hours (hr). Activation of RIPK1 was assessed by S166 phosphorylation by Western blot (p-RIPK1). Positive controls were MDA-MB-231 cells treated with 10nM of birinapant and 25uM Z-VAD-FMK for 12 hr. The ratio of p-RIPK1/RIPK1 was calculated using densitometry. **(b)** TNBC R- (orange) and S- (blue) PDxOs were treated with birinapant (10nM) and Z-VAD-FMK (25uM, 50uM) +/- the RIPK1 kinase inhibitor Nec-1s (25uM) for 24 hours and analyzed as described in panel **a**. **(c)** TNBC S-PDxOs were pre-treated overnight with Nec-1s followed by combined titration of Nec-1s and birinapant at the indicated concentrations for an additional 48 hours. Cell viability after treatment was evaluated with Cell Titer Glo (3 biological replicates with 4 technical replicates each). For each biological replicate, the area under the curve (AUC) for each combination treatment row was compared with the corresponding AUC for birinapant alone as the reference control (row 0; ref = reference) to determine the effects of Nec-1s on birinapant response. Statistical significance between AUCs was assessed using a two-sided paired Student’s *t*-test (n = 3 biological replicates), and p-values are color-coded by significance. **(d)** TNBC S-PDxOs were pre-treated for 12 hours with LD4172 (100nM) or vehicle, then treated with birinapant (10nM) +/- LD4172, in the presence of Z-VAD-FMK (25uM) for an additional 12 hours. Activation of RIPK1 was assessed by S166 phosphorylation via Western blot (p-RIPK1). **(e)** TNBC S-PDxOs were pre-treated overnight with titrated concentrations of the RIPK1 degrader LD4172 or vehicle, and treated with titrated LD4172 or vehicle, plus Birinapant (16nM), at the indicated concentrations for an additional 48 hours. Cell viability was determined as in panel **c** (3 biological replicates with 4 technical replicates each). For all heatmaps, darker blue indicates cytotoxicity, white indicates a cytostatic response, yellow indicates growth, and red p-values indicate statistical significance. Detailed statistics for single-agent heat map data and calculated AUC values for each biological replicate are shown in **Supp. Fig. 4**.

To determine if RIPK1 autophosphorylation was required for birinapant-induced death, we pretreated PDxOs with Nec-1s, a selective RIPK1 kinase inhibitor^9,27^, and continued Nec-1s treatment in the presence of birinapant. Nec-1s blocked birinapant-induced RIPK1 S166 autophosphorylation in S-PDxOs (**Fig. 3b**), and reduced birinapant-mediated cell death, as assessed by cell viability (**Fig. 3c**) and comparing areas under the dose response curve (AUCs; **Supp. Fig. 4a**; vehicle controls shown in **Supp. Fig. 3c**). As secondary confirmation of the requirement for RIPK1 for cell death following birinapant treatment, we pretreated S-PDxOs with the RIPK1 protein degrader LD4172^28^ and continued treatment in the presence of birinapant. We observed degradation of RIPK1 in the presence of LD4172, and decreased RIPK1 autophosphorylation during birinapant and LD4172 co-treatment (**Fig. 3d**). Degradation of RIPK1 protein blocked birinapant-induced cell death (**Fig. 3e**, statistics shown in **Supp. Fig. 4b**). These data demonstrated that RIPK1 protein and kinase activity were required for cell death following birinapant treatment in S-PDxOs.

**Figure 4.**
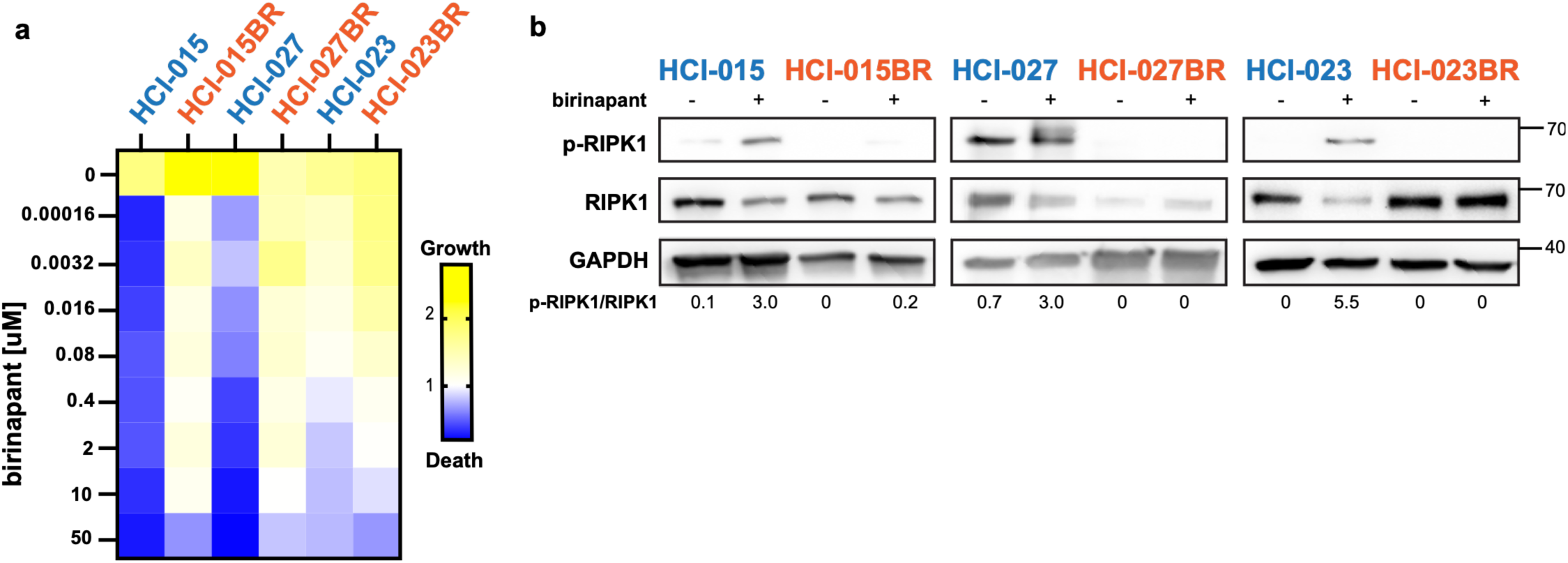
Acquired birinapant resistance is associated with failure to activate RIPK1. **(a)** Response of parental PDxOs (blue) and PDxO sublines derived from PDX selected for acquired birinapant resistance (BR, orange) to 6 days of birinapant treatment at the indicated concentrations. Cell viability was evaluated with Cell Titer Glo (3 biological replicates with 4 technical replicates each). Darker blue indicates cytotoxicity, white indicates a cytostatic response, and yellow indicates growth. **(b)** TNBC R- (orange) and S- (blue) PDxOs were treated with birinapant (10nM) in the presence of Z-VAD-FMK (25uM) for 12 hours. Activation of RIPK1 was assessed by S166 phosphorylation via Western blot (p-RIPK1). The ratio of p-RIPK1/RIPK1 was calculated using densitometry. Detailed statistics for the heat map data are shown in **Supp. Fig. 5b-c**.

Based on our data indicating that failure of RIPK1 autophosphorylation was associated with *de novo* birinapant resistance in R-PDxOs, we asked if lack of RIPK1 activity was also associated with acquired resistance to birinapant. We developed PDX models of acquired resistance using an intermittent birinapant dosing strategy in three different S-PDX lines. When tumors became resistant (‘BR’ lines; **Supp. Fig. 5a**), we established PDxOs^29^ from the resistant tumors. All three BR PDxO lines showed resistance to birinapant in vitro compared to their parental line (**Fig. 4a**, statistics shown in **Supp. Fig. 5b-c**), and they each displayed little or no RIPK1 autophosphorylation following birinapant treatment compared to the parental S-PDxOs (**Fig. 4b**). One of the lines, HCI-27BR, also showed notable reduction of total RIPK1 protein with acquired resistance, consistent with our prior result that eliminating RIPK1 protein was sufficient to facilitate birinapant resistance (**Fig. 3d**). These data demonstrated that defective RIPK1 activation is associated with both *de novo* and acquired resistance to birinapant in R-PDxOs.

**Figure 5:**
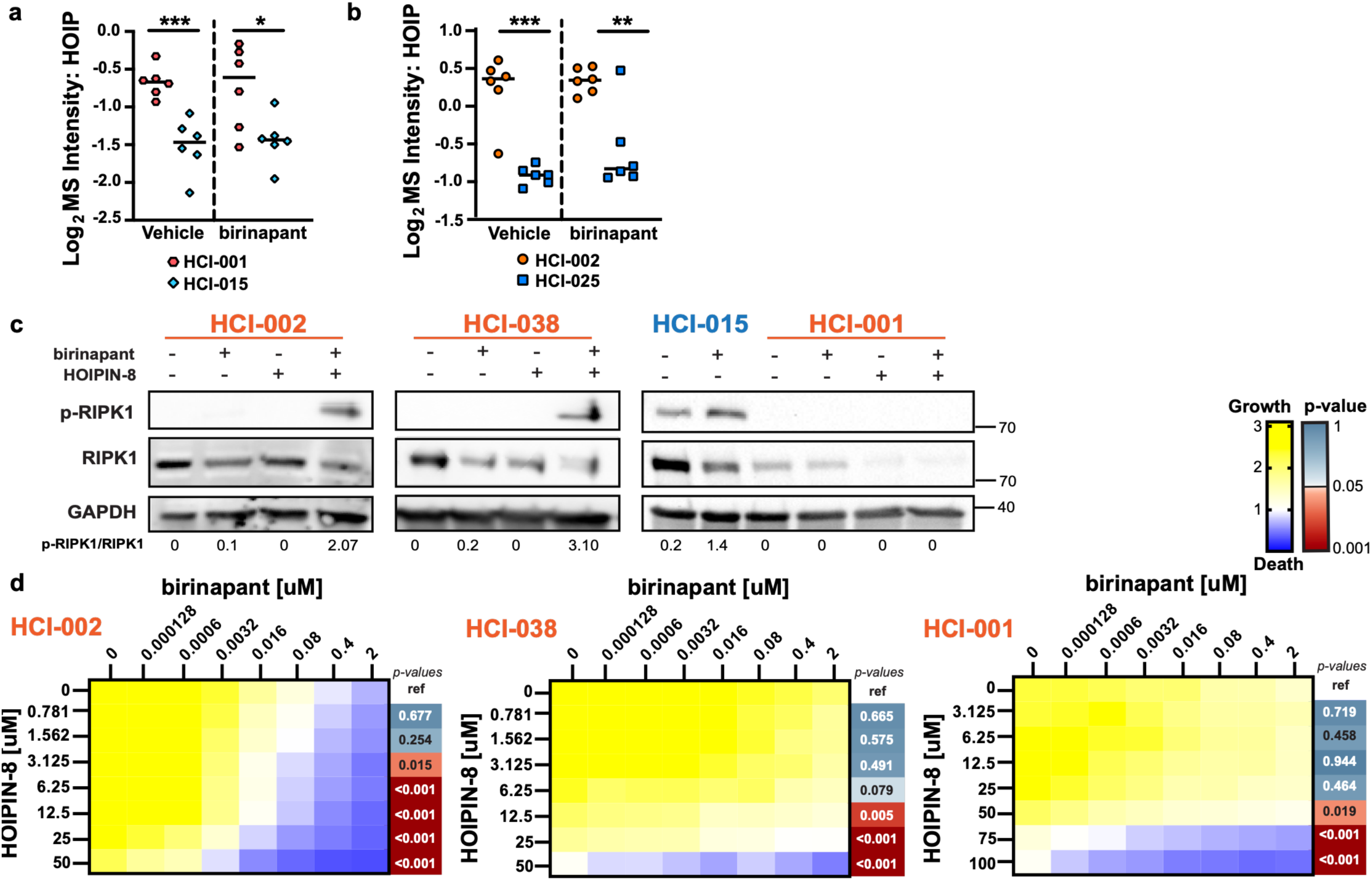
HOIP inhibition restores RIPK1 activation and sensitizes resistant TNBC PDxOs to birinapant (a-b) TNBC R- (orange) and S- (blue) PDxOs were treated with vehicle or birinapant (10nM) in the presence of Z-VAD-FMK (25uM) for 12 hours (n=6 replicates per condition). Protein was extracted and two independent LC-MS/MS analyses were performed. HOIP fold change was calculated within each batch (a, HCI-001 vs HCI-015; b, HCI-002 vs HCI-025) to mitigate batch effects. Statistical significance was calculated using a two-sided unpaired Student’s t-test (HCI-001 vs HCI-015: **p=0.007,***p=0.001; HCI-002 vs HCI-025 *p=0.029, **p=0.001,) **(c)** TNBC R-PDxOs were treated with birinapant (10nM), HOIPIN-8 (HCI-002, HCI-038: 50uM, HCI-001: 75uM; concentrations selected to induce cell death from panel **d**), or the combination in the presence of Z-VAD-FMK (25uM) for 12 hours. Conditions that contained HOIPIN-8 were pre-treated for 12 hours before adding birinapant and Z-VAD-FMK. The S-PDxO HCI-015 was used as a positive control. Activation of RIPK1 via S166 phosphorylation was assessed by Western blot (p-RIPK1). The ratio of p-RIPK1/RIPK1 was calculated with densitometry. **(d)** TNBC R-PDxOs were pre-treated overnight with HOIPIN-8, then treated with HOIPIN-8 and birinapant at the indicated concentrations for an additional 48 hours. Cell viability after treatment was evaluated with Cell Titer Glo (3 biological replicates with 4 technical replicates each). For each biological replicate, the area under the curve (AUC) for each combination treatment row was calculated and compared with the corresponding AUC for birinapant alone (row 0; ref = reference) to determine the effects of HOIPIN-8 on birinapant response. Statistical significance between AUCs was assessed using a two-sided paired Student’s *t*-test (n = 3 biological replicates). For all heatmaps, darker blue indicates cytotoxicity, white indicates a cytostatic response, and yellow indicates growth, and red p-values indicate statistical significance. Calculated AUC values are shown in **Supp. Fig. 6**.

### HOIP regulation of RIPK1 autophosphorylation contributes to birinapant resistance

We next sought to investigate the molecular mechanism impeding RIPK1 autophosphorylation in R-PDxOs following birinapant treatment. To identify candidate proteins that could be contributing to this resistance phenotype, we performed two independent LC-MS/MS global proteomics experiments comparing R- and S-PDxOs treated with vehicle or birinapant (HCI-001 vs HCI-015 and HCI-002 vs HCI-025) and examined proteins involved in TNFR signaling. We discovered that, in both experiments, the E3 ubiquitin ligase HOIP was the only TNFR-pathway associated protein that was more abundant in R-PDxOs compared to S-PDxOs, both at baseline and after birinapant treatment (**Fig. 5a-b**). HOIP, along with HOIL-1L and SHARPIN, is recruited to RIPK1 following its K63 ubiquitination by cIAPs, forming the linear ubiquitin chain assembly complex (LUBAC) and adding M1 linear ubiquitin chains to RIPK1^27,30–35^. These ubiquitin chains tether RIPK1 to the TNFR, stabilizing the NF-κB signaling complex, and result in pro-inflammatory survival signaling^36^. M1 ubiquitination has been reported to inhibit RIPK1 autophosphorylation and TNFR cell death signaling under certain inflammatory conditions^12,32^ but has not been reported to promote resistance to SMAC mimetic treatment in tumors.

To determine if we could restore RIPK1 activation and subsequent sensitivity to birinapant by inhibiting HOIP ubiquitin ligase activity, we treated R-PDxOs with the HOIP-targeted LUBAC inhibitor HOIPIN-8^31,37,38^ and tested RIPK1 phosphorylation and cell viability following birinapant treatment. Indeed, RIPK1 activation was restored by the addition of HOIPIN-8 during birinapant treatment in two out of three R-PDxOs tested (HCI-002 and HCI-038; **Fig. 5c**). Two of the lines, HCI-001 and HCI-038, had reduced total RIPK1 in the presence of HOIPIN-8 (**Fig. 5c**). HOIPIN-8 significantly sensitized all three R-PDxOs to birinapant-mediated death, albeit to varying degrees (**Fig. 5d**; AUCs in **Supp. Fig. 6b**; vehicle controls in **Supp. Fig. 6c**). Interestingly, in R-PDxOs in which birinapant resistance could be reversed by HOIPIN-8, birinapant treatment increased M1 ubiquitination, and HOIP inhibition reduced it back to baseline levels (**Supp. Fig. 6a**). Together, these data demonstrated that restoration of RIPK1 autophosphorylation by blocking HOIP/LUBAC activity was sufficient to resensitize some R-PDxOs to birinapant, indicating that RIPK1 activity plays a crucial role in both birinapant response and resistance in TNBC.

## Discussion

This study investigated the molecular basis of birinapant resistance in patient-derived models of TNBC and identified defects in RIPK1 activation as a key determinant of resistance. We discovered that in both *de novo* and acquired resistant TNBC PDxOs, birinapant failed to induce RIPK1 autophosphorylation, an established step in SMAC mimetic-induced cell death^9,14,39,40^. Consistent with the requirement for RIPK1 activity, pharmacologic inhibition of RIPK1 kinase activity or depletion of RIPK1 protein protected S-PDxOs from birinapant. On the other hand, inhibiting HOIP, the E3 ubiquitin ligase component of the LUBAC complex^31,33^, restored birinapant sensitivity in R-PDxOs, with concomitant restoration of RIPK1 activity in two out of three cases. These data provide new insight supporting a model in which persistent LUBAC activity during birinapant treatment prevents RIPK1 autophosphorylation and downstream cell death signaling, promoting resistance during treatment.

Alternative mechanisms of SMAC mimetic resistance have been described, primarily centering on persisting IAPs or disrupted complex IIb formation. Compensatory upregulation of cIAP2 during SMAC mimetic treatment was observed in lung cancer cell lines and mouse embryonic fibroblasts (MEFs), conferring resistance^41,42^. In our models, however, IAPs were degraded to comparable levels in S- and R-PDxOs, indicating that incomplete target degradation does not explain resistance. Failure to form complex IIb despite cIAP2 degradation during SMAC mimetic treatment has been demonstrated in melanoma, leukemia, and oral squamous cell carcinoma cell lines^43–46^. It was also noted that persisting expression of c-FLIP, an anti-apoptotic regulator that inhibits complex IIb formation^47^, could promote resistance^43,46,47^. In one instance, melanoma cells exhibited RIPK1 activation during birinapant treatment, but remained resistant until c-FLIP protein levels were reduced^46^. Interestingly, HOIP/LUBAC has been identified as a mediator of c-FLIP stability; HOIP knockdown resulted in c-FLIP degradation, and increased sensitivity to SMAC mimetic treatment^48^. Because c-FLIP acts downstream of RIPK1 activation at complex IIb to inhibit apoptosis, we postulate that deficient RIPK1 autophosphorylation remains the primary driver of birinapant resistance in TNBC. However, it is plausible that HOIP inhibition in R-PDxOs may promote c-FLIP degradation in addition to RIPK1 activation, thereby enabling downstream apoptotic signaling and further sensitizing these R-PDxOs to birinapant. Future work should determine if there is a relationship between RIPK1 activity and c-FLIP abundance during HOIP inhibition, and if that contributes to birinapant sensitization in TNBC.

Our results also raise mechanistic questions about how HOIP restrains RIPK1 activation during birinapant treatment. The finding that birinapant stimulates the activity of HOIP/LUBAC-dependent M1 ubiquitination, which is known to block RIPK1 activity^30,36,49^, supports our findings that dysregulation of RIPK1 activity is a key mediator of birinapant resistance. However, this result was surprising, as cIAPs facilitate recruitment of the LUBAC complex to the TNFR to regulate RIPK1 activity through M1 ubiquitin^26,29^. Since cIAP1/2 were degraded with birinapant treatment in our PDxOs, the observed global increase in M1 ubiquitin may represent an adaptive response that preserves pro-survival signaling and supports continued tumor-cell proliferation^27,30,50,51^. Further investigation into the relationship between cIAP degradation and LUBAC activity is needed to elucidate this mechanism. Second, M1 ubiquitination of RIPK1 by HOIP has been shown to recruit TBK1, IKKα/β, and TAK1 kinases to the TNFR, facilitating phosphorylation of RIPK1 at inhibitory sites and preventing its autophosphorylation and dissociation from the TNFR^12,52–56^. Increased M1 ubiquitination of RIPK1 at the TNFR during birinapant treatment in R-PDxOs would be predicted to suppress RIPK1 autophosphorylation through recruitment of these inhibitory kinases. Testing kinase recruitment and site-specific inhibitory phosphorylation after birinapant treatment, with and without HOIP inhibition, will be necessary to evaluate this mechanism.

HOIP may also influence RIPK1 protein stability in some cases. In two R-PDxOs, we observed that HOIP inhibition reduced total RIPK1 abundance. One potential explanation for this loss of RIPK1 protein in the presence of HOIPIN-8 is that HOIP/LUBAC-dependent M1 ubiquitination may protect RIPK1 from destabilization, as has been shown with another apoptosis regulator, c-FLIP^48^. Knockdown of HOIP reduced M1 ubiquitination of c-FLIP, which permitted increased K48-linked ubiquitination and proteasomal degradation of c-FLIP^48^. Whether HOIP/LUBAC inhibition promotes degradative K48-linked ubiquitination and degradation of RIPK1 in R-PDxOs is unknown. We also observed that one PDxO model exhibited loss of RIPK1 protein during HOIP inhibition and failed to induce RIPK1 autophosphorylation upon birinapant co-treatment. Because our depletion experiments showed that RIPK1 protein is required for birinapant-induced death in S-PDxOs, this loss of RIPK1 could also limit the ability of HOIP inhibition to restore sensitivity. In this same line, the cell death observed with birinapant at high HOIPIN-8 concentrations might reflect a RIPK1-independent mechanism of cell death. Therefore, determining the prevalence of HOIPIN-8–mediated RIPK1 loss in a larger cohort of R-PDxOs and assessing whether these models can be resensitized to combined HOIPIN-8 and birinapant treatment represent important future directions.

There are limitations of this study that define important directions for future work. First, although caspase-3 cleavage and reduced ATP content demonstrate caspase-associated cell death, they do not distinguish apoptosis from pyroptosis. These pathways share several features downstream of RIPK1, including cleavage of caspases 8, 3, and 7^57^. Therefore, additional pathway-specific markers and perturbation experiments are required to exclude pyroptosis as a potential mode of birinapant-induced cell death. Second, our experiments establish a requirement for RIPK1 activity in birinapant-induced cell death but do not demonstrate that signaling is TNF-α or TNFR-dependent. Birinapant could also promote formation of the ripoptosome, a death-signaling complex similar to complex IIb that forms after IAP degradation, independently of TNFR activation^58^. Assessing TNF/TNFR dependence using a TNF-α-neutralizing antibody and directly measuring formation of complex IIb and the ripoptosome during birinapant treatment will clarify exactly how RIPK1 functions in S-PDxOs and may reveal additional mechanisms by which some TNBCs evade cell death following birinapant treatment.

Together, these findings identify defective RIPK1 activation as a mechanism of birinapant resistance in TNBC PDxOs and implicate persistent HOIP-mediated M1 ubiquitination as one means of enforcing this impaired state. This improved understanding of the requirements for efficacy of SMAC mimetics like birinapant provides a framework for refining biomarkers for response to this class of therapeutics and for developing strategies to reverse resistance in selected TNBC tumors.

## Methods

### Cell culture

Stable PDxO lines were established from patient-derived xenografts (PDXs) and cultured as previously described^18,29^. Briefly, PDxOs were maintained as embedded cultures in growth factor-reduced Matrigel (Corning) domes and complete medium (Advanced Dulbecco’s Modified Eagle Medium/Ham’s F-12 (DMEM/F-12) (ThermoFisher), with 5% fetal bovine serum (FBS), 10mM HEPES, 1x Glutamax, 1ug/mL Hydrocortisone, 50ug/mL Gentamycin, and 10ng/mL human epidermal growth factor (EGF)), supplemented with 10uM Y-27632 was added fresh at the time of medium exchange or passaging. MDA-MB-231 cells were obtained from American Type Culture Collection (ATCC) and maintained in complete medium (DMEM + Glutamax (ThermoFisher) with 10% FBS, 1% Penicillin-Streptomycin, and 1% Sodium Pyruvate). Cell cultures and organoids were maintained using standard tissue culture practices and housed in a humidified incubator at 37°C and 5% CO_2_. For drug treatment conditions evaluated via Western blot or qRT-PCR, cells were treated in their normal culture conditions with the indicated drugs diluted to their final concentrations in complete culture medium, for 6-24 hours (harvest timepoints indicated in figure panels). PDxOs in long-term culture tested negative for mycoplasma.

### In vivo experiments

All procedures were approved by the University of Utah’s Institutional Animal Care and Use Committee. Five- to six-week-old female NOD.Cg-*Rag1*^tm1Mom^*Il2rg*^tm1Wjl^/SzJ (NRG) mice (The Jackson Laboratory, stock no. 007799) were orthotopically implanted with thawed human breast cancer PDX fragments into the cleared inguinal mammary fat pad, as previously described^59^. Tumor growth was monitored weekly using digital calipers. To select birinapant-resistant sublines, mice received birinapant (20 mg/kg; obtained previously National Cancer Institute CTEP, in 15% captisol in saline) by intraperitoneal injection three times per week, starting when tumors reached approximately 100 mm³. During treatment, tumors were measured twice weekly. Treatment was discontinued after 35 days, and monitoring continued off-treatment until tumors regrew to approximately 100 mm³. Birinapant treatment was then resumed using the same dose schedule until tumor volume reached 500 mm³ (endpoint criteria), at which time tumors were harvested. For caspase assays, mice bearing tumors of approximately 100 mm^3^ were given a single dose of birinapant as described above, and tumors were harvested at the indicated time points. Tumors were prepared as specified previously^60^ to extract protein. Cleaved caspase-3 was measured via Luminex bead multiplex immunoassay and analyzed as previously described^60^.

### Western blots and analysis

Cells were harvested and lysed in radioimmunoprecipitation assay (RIPA) buffer (Tris-HCL PH 7.4 [50mM], Igepal NP-40 [1x], NA-deoxycholate [0.5x], SDS [0.1%], NaCl [150mM], EDTA [5mM]), with protease (Sigma, P8340) and phosphatase (Sigma, P5726 & P0044) inhibitor cocktails (1x), β-glycerophosphate (1mM), sodium fluoride (5mM), PMSF (5mM), sodium pyrophosphate (10mM), dithiothreitol (DTT) (1mM), plus fresh N-Ethylamaleimide (20mM) for M1 Ubiquitin studies. Protein concentration was quantified using the DC Protein Assay Kit (Bio-Rad, 500-0116). Samples were mixed with 4X Laemmeli Buffer (Bio-Rad 1610747) containing 1mM DTT and boiled at 95°F for 5 minutes to denature. After separation via SDS-PAGE, proteins were transferred to a polyvinylidene difluoride (PVDF) membrane (Millipore Sigma) using a wet transfer. Membranes were blocked with 5% bovine serum albumin (BSA) with 0.1% TWEEN 20 (Sigma) and probed with their respective primary antibodies overnight at 4°C. Primary antibodies were diluted in 5% BSA in TBST at 1:1000, and secondary antibodies were diluted in 5% BSA at 1:10,000. Membranes were developed using Clarity Western ECL Substrate (Bio-Rad) and images were captured using the Azure Biosystems 400 Western Blot Imager or the Bio-Rad ChemiDoc MP. Results were analyzed via densitometry using Fiji^61^ (ImageJ), with ratios indicated in the figure panel. RIPK1 activity was assessed by calculating the ratio of p-RIPK1 to total full-length RIPK1.

### Antibodies and other reagents

The following antibodies were used for Western Blotting: GAPDH XP (Cell Signaling Technology, 5174; 1:2000), β-actin (Cell Signaling Technology, 1470; 1:1000), RIPK1 (Cell Signaling Technology, 3493; 1:1000), p-RIPK1 S166 (Cell Signaling Technology, 657346; 1:1000), cIAP1 (Cell Signaling Technology, 7065; 1:1000), cIAP2 (Cell Signaling Technology, 3130; 1:1000), XIAP (Cell Signaling Technology, 2045; 1:1000), IκB (Cell Signaling Technology, 4814; 1:1000), p-IκB (Cell Signaling Technology, 2859;1:1000), TRADD (Cell Signaling Technology, 3684; 1:1000), TRAF2 (Cell Signaling Technology, 4712; 1:1000), Caspase-8 (Cell Signaling Technology, 9746; 1:1000), FADD (Cell Signaling Technology, 2782; 1:1000), TNFR1 (Cell Signaling Technology, 3736; 1:1000), M1 Ubiquitin (Millipore Sigma, MABS451; 1:1000), HOIP (Cell Signaling Technology, 99633; 1:1000), Goat α-Rabbit Secondary Antibody (Invitrogen, 31460; 1:10,000), Goat α-Mouse secondary antibody (Invitrogen, A16078; 1:10,000). The following reagents were used for treatments in culture: human recombinant TNF-alpha (R&D Systems, 210-TA), human recombinant TRAIL/TNFSF10 (R&D Systems, 375-TL), staurosporine (Selleck Chemicals, S1421), birinapant (MedChem Express, HY-16591), ASTX660 (Selleck Chemicals, S8681), GDC0152 (MedChem Express, HY-13638), GDC0917 (MedChem Express, HY-15835), LCL161 (Selleck Chemicals, S7009), Z-VAD-FMK (Selleck Chemicals, S7023), Necrostatin-1s (Selleck Chemicals, S8641), HOIPIN-8 (MedChem Express, HY-122882).

### PDxO Cell Titer Glo assays

Seeding, treatment, and endpoint collection of PDxOs treated with the indicated drugs was performed as described previously^18^. Briefly, mature PDxOs were harvested from factor-reduced Matrigel (Corning) (Corning, 354230) using dispase II (Themo Fisher) treatment for 25 minutes at 37°C. Approximately 50 organoids were seeded in each well of a 384-well opaque tissue culture plate containing a 10uL Matrigel base layer, with an additional 16 wells seeded in a separate 384-well plate as an untreated baseline control plate to be collected on day 1. For every biological replicate, PDxOs were seeded in technical quadruplicate, in 30uL of complete media supplemented with fresh 10uM Y-27632. If indicated that PDxOs were pre-treated overnight, media contained the indicated drugs. Plates were incubated overnight at 37°C and 5% CO_2_. Thirty microliters of complete medium was added to the control plate, which was assayed using the CTG-3D reagent to assess ATP content as a surrogate for viability, which served as the day 0 seeding baseline controls. Thirty microliters of complete media containing the indicated drugs was added to the treatment plate(s) which were sealed using Breathe-Easy sealing membranes (Millipore Sigma) and incubated for either 48hr (mechanistic studies) or 6 days (**Fig. 1**, overall viability screening) at 37°C and 5% CO_2_. Dosed plates were assayed using the CTG-3D reagent to assess ATP content as a surrogate for viability. Luminescent values for each technical quadruplicate were normalized to their average day 0 control values and each set of normalized quadruplicates was averaged together to obtain one value per biological replicate. For each experiment, biological replicates (n=3) were averaged, and growth during treatment is shown in yellow, cytostatic response (no growth) in white, and cytotoxic response (death) in blue.

### RNA extraction and RT-qPCR

RNA from PDxOs was collected and analyzed via RT-qPCR using methods previously described^18^. Briefly, mature PDxOs were collected and lysed in Buffer RLT Plus (Qiagen) containing 1% β-mercaptoethanol (Sigma-Aldrich) and stored at -80°C before being rapidly thawed at 37°C for 5 minutes and centrifuged using a QiaShredder (Qiagen). RNA was isolated using the AllPrep DNA/RNA Mini Kit (Qiagen) and quantified using the Qubit RNA Broad Range assay kit (Invitrogen), following the manufacturer’s instructions. cDNA was prepared (SuperScript IV VILO MasterMix with ezDNase Enzyme kit (Invitrogen)) and 1ng of cDNA was used in each reaction, with 4 technical replicates per condition, using the PowerUp SYBR Green Master Mix (Applied Biosystems) and PrimeTime qPCR primers (IDT, RIPK1 forward: 5’-ACC CAT CCC AAG TCC CT-3’, reverse: 5’-CTT GAT TGT GTG GCA GTA CCT-3’; GAPDH forward: 5’-TGT AGT TGA GGT CAA TGA AGG-3’, reverse: 5’-ACA TCG CTC AGA CAC CAT-3’). Quantitative PCR was performed using a Roche LightCycler 480. Relative gene expression was normalized to GAPDH and fold change was calculated using the 2^−ΔΔCt^ method^62^.

### Global proteome profiling

Global proteome analysis by nLC-MS was performed as described previously^63^. Briefly, PDxOs were lysed by adding 750 µL of 8 M urea lysis buffer also containing 100 mM Tris (pH = 8.5), 5 mM Tris(2-carboxyethyl)phosphine (TCEP) and 10 mM chloroacetamide (CAM) and incubation on an end-over-end rotator for 30 min at room temperature. Lysates were then clarified by centrifugation at 21,000 rcf at RT, and protein concentration was determined using the Pierce 660 nm assay. Aliquots of 100 µg of protein in lysis buffer were pipetted into a new microtube and diluted with four times the volume ice-cold acetone. 80% aq. trichloroacetic acid (TCA) was added to a final concentration of 4% (v/v), samples were vortexed briefly at max speed and kept at -20°C overnight. Precipitated protein was pelleted at 2,000 rcf at 4°C for 10 min and the supernatant aspirated. The pellet was cleaned by adding 1 mL of ice-cold acetone, dispersing the pellet in a sonicator bath, pelleting the protein at 2,000 rcf at 4°C for 10 min, and aspirating the supernatant; this step was repeated once more. The pellets were dried until translucent (2-3 min), 100 µL of 8 M urea buffer containing 100 mM Tris (pH = 8.5) was added and the sample agitated on a thermal shaker at 1,400 rpm at 37°C until the pellet was dissolved (15-30 min). The samples were diluted 2-fold with 100 mM triethylammonium bicarbonate (TEAB), the pH was adjusted to 8-9 with 1 M aq. NaOH, if needed, and 1 µg of Endproteinase Lys-C (Wako-Fujifilm) was added (1:100 ratio digestive enzyme to protein). Then, the mixtures were agitated on a thermal shaker at 1,400 rpm at 37°C for 2 h, diluted another 2-fold with 100 mM TEAB, and 1 µg of MS grade trypsin was added (Pierce). The pH was tested and readjusted if needed. The mixtures were agitated on a thermomixer at 1,400 rpm at 37°C overnight, acidified with formic acid (FA, 1% final), and cleared by centrifugation for 10 min at 14,000 rcf at RT. An aliquot of the supernatant equaling 5 µg peptide was desalted on C18 StageTips to be used for global proteome analysis by nLC-MS.^64^

### nLC/MS analyses of peptide samples

Peptide samples of 200 ng were analyzed on a nanoElute 2 – timsTOF Pro 2 nLC-MS system (Bruker) as described^63^. Briefly, peptides were separated using 15 cm long, 150 µm inner diameter PepSep columns packed with 1.5 µm diameter C_18_ beads (Bruker) in 45 min long LC gradients (5-30% B) at flow rates of 0.5 µL/min. LC solvents were A (0.1% formic acid in LC-MS-grade water) and B (0.1% formic acid in LC-MS-grade acetonitrile). MS data were acquired in data-independent acquisition mode using the diaPASEF method^65^. General MS instrument parameters were the following: polarity: positive; mass range: 100 m/z to 1700 m/z; 1/K_0_ start: 0.60 Vs/cm²; 1/K_0_ end: 1.40 Vs/cm², CaptiveSpray; capillary: 1600 V.

### Computation of Bruker MS raw files and output data processing

Computation of Bruker MS raw files and output data processing was performed as described^63^. Briefly, Bruker MS raw files were computed using DIA-NN v.1.8.1 in library-free mode^66^. The Homo sapiens FASTA downloaded from UniProt on 07.14.2023 (UP000005640) was used in the DIA-NN search. DIA-NN search parameters were the following: reannotate: enabled; FASTA digest for library-free search/library generation: enabled; deep learning-based spectra, RTs, and IMs prediction: enabled; protease: Trypsin/P; missed cleavages: 1; number of variable modifications: 0; N-term M excision: enabled; C carbamidomethylation: enabled; peptide length range: 7 - 30; precursor charge range: 1 - 4; precursor m/z range: 300 - 1800; fragment ion m/z range: 200 – 1800; generate spectral library: enabled; quantities matrices: enabled; precursor FDR (%): 1.0; mass accuracy: 0.0; MS1 accuracy: 0.0; scan window: 0; use isotopologues: enabled; MBR: enabled; no shared spectra: enabled; protein inference: Genes; neural network classifier: Single pass mode; quantification strategy: Robust LC (high precision); cross-run normalization: global; smart profiling; speed and RAM usage: optimal results. DIA-NN raw output files were loaded into Perseus v2.0.10.0, log2 transformed, median-normalized, and missing values were imputed (width: 0.2; downshift 1.8)^67^. Perseus was also used for differential expression analysis.

### Statistical analyses

All statistical tests for viability assays were performed using the averaged calculated Cell Titer Glo growth values for three biological replicates (n=3), which each comprised technical quadruplicates and are displayed as detailed in the Cell Titer Glo methods above. Statistical analysis of normalized growth between each treatment condition and its plate-matched reference were evaluated and visualized as *P*-value heat maps. Reference (control) conditions were defined according to the experimental setting: (1) for single-agent experiments, each drug concentration was compared with the untreated condition (top row of each heat map, denoted as “ref”) for the same PDxO line; (2) for combination treatment experiments, the calculated area under the curve (AUC) values (n=3) for each row were compared with the corresponding condition of birinapant alone (top row of each heat map, denoted as “ref”). Because variance estimates based on three biological replicates are inherently imprecise, variance was pooled across the *k* treatment conditions within each predefined comparison group. Pooling groups comprised the concentrations tested within an individual PDxO line; the rows within a treatment arm, or the combination agent rows. For each condition, the test statistic was calculated as t = (mean difference) / √(pooled variance / 3), with 2k degrees of freedom, two-sided. Variance was never pooled across PDxO lines or across treatment arms.

## Data & Resources Availability

Raw Bruker mass spectrometry files, DIA-NN output files and all associated metadata have been deposited in the MassIVE repository (University of California, San Diego) under accession number MSV000103621.

## Author Contribution Statement

E.B., P.J., A.S., R.P., and J.D performed experiments. C.Y., and E.C.S. performed the drug screening shown in Fig. 1. Z.C. performed all in vivo experimentation. C.Y. and Y.Q. performed statistical analysis. M.G. performed MS analysis and statistics. J.F. and A.W. supervised the work. E.B., J.F., and A.W. wrote the manuscript. All authors reviewed and edited the manuscript.

## Acknowledgements

This work was conducted with funding from the National Cancer Institute (U54CA224076 to A.L.W., B.E.W, and the Huntsman Cancer Institute Cancer Genetics, Epigenetics, Models, & Signaling training grant T32CA265782 supporting E.B). Research reported in this publication utilized the Preclinical Cancer Models Shared Resource at Huntsman Cancer Institute at the University of Utah, supported by the National Cancer Institute of the National Institutes of Health under Award Number P30CA042014. The content is solely the responsibility of the authors and does not necessarily represent the official views of the NIH. We acknowledge the use of OpenAI’s ChatGPT (GPT-5) to improve the clarity and readability of this manuscript. All AI-assisted content was reviewed and edited by the authors, who assume full responsibility for the accuracy and integrity of the final manuscript.

---

**Supplementary Figure 1:**
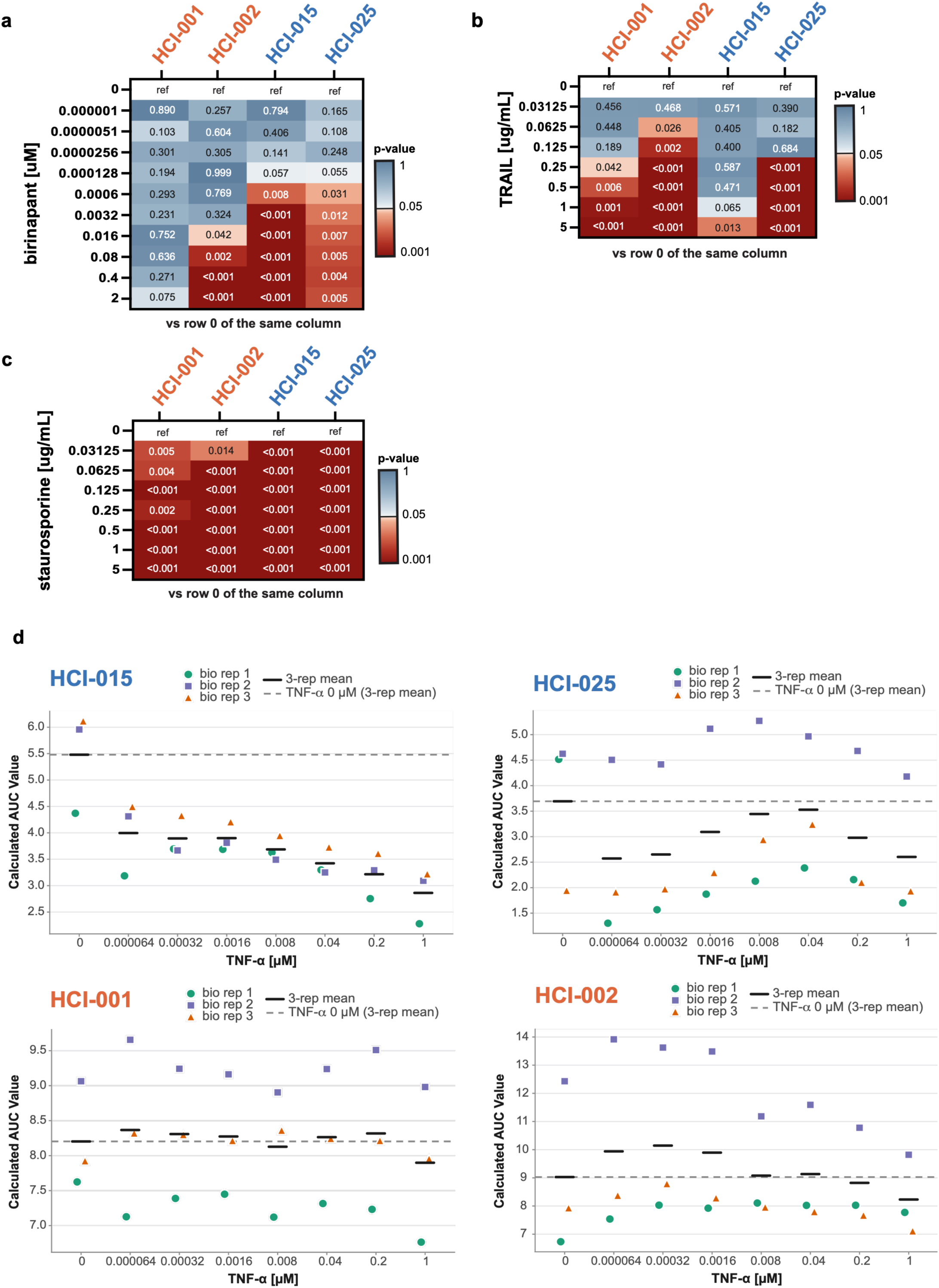
Birinapant-resistant PDxOs are sensitive to TRAIL and staurosporine-mediated cell death, but not to TNFR-mediated cell death. (a-c) P-value heatmaps corresponding to the drug response heatmaps shown in **Fig. 2d-f** (R-PDxOs in orange, S-PDxOs in blue). For each biological replicate, viability at each treatment concentration was compared to the corresponding non-treated control (row 0; ref=reference). Statistical significance was assessed using a two-sided paired Student’s *t*-test (*n* = 3 biological replicates), and p-values are color-coded by significance. **(d)** Area under the curve (AUC) values calculated corresponding to the drug response heatmap in **Fig. 2g** to depict variance between biological replicates (bio rep, n = 3). The AUC replicate dot plots show the calculated AUCs at each concentration of TNF-α treatment (μM; x-axis) to evaluate of the effects of TNF-α addition on the birinapant dose-response curve. (solid black line = mean; dashed line = AUC of row 0). Because biological replicates are from different PDxO passages, there is inherent variability in growth rate that influences the ATP content (viability) over the duration of the assay relative to the Day 0 control, influencing the calculated AUC.

**Supplementary Figure 2:**
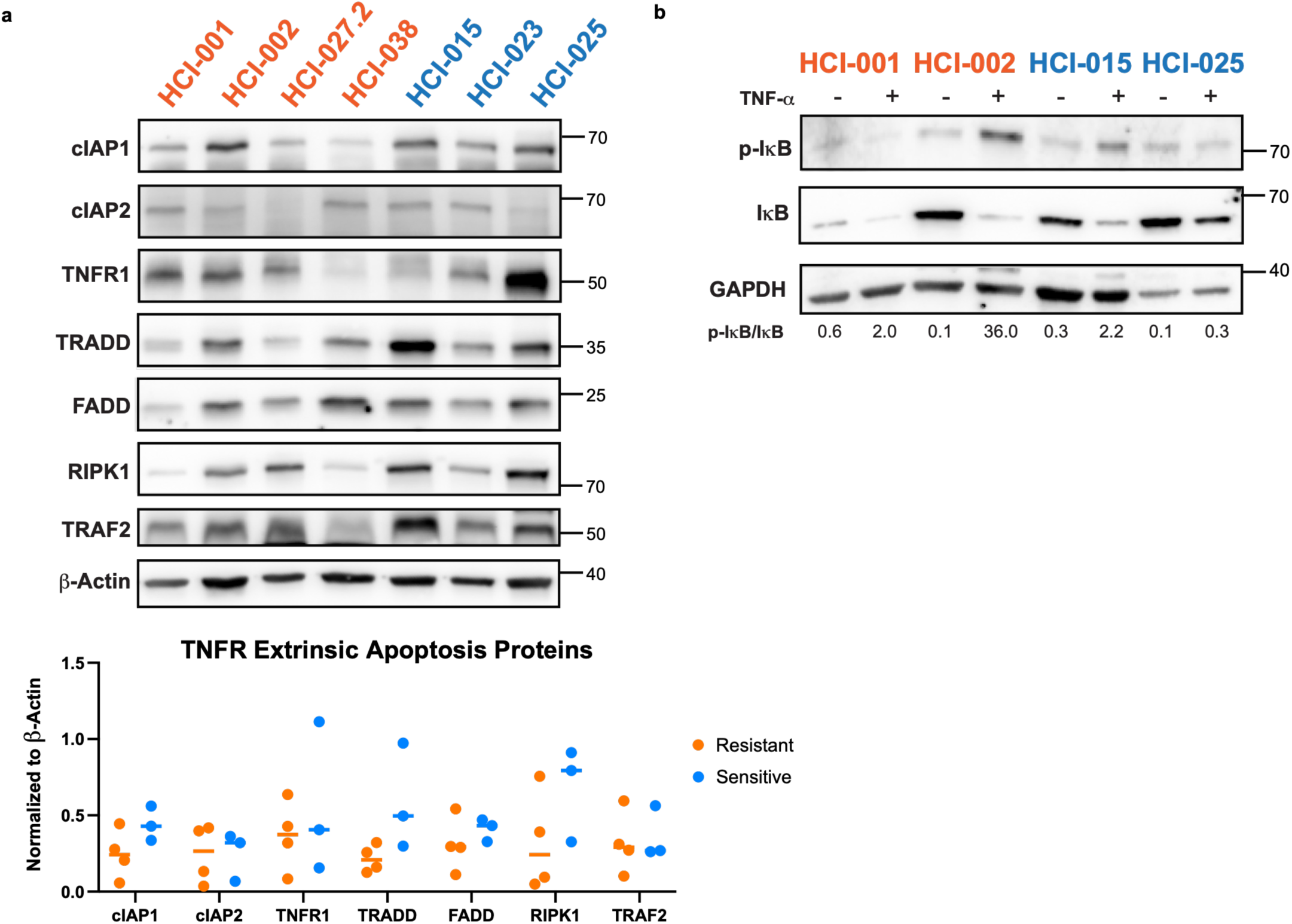
Both resistant and sensitive TNBC PDxOs express proteins required for TNFR-mediated signaling. **(a)** Baseline expression levels of components of the TNFR signaling pathway were assessed in R- (orange) and S- (blue) TNBC PDxOs by Western blot (top). Quantification was performed using densitometry, and protein abundance was normalized to β-Actin (bottom). **(b)** R- and S-PDxOs were treated with TNF-α (20 ng/mL) for 30 minutes to evaluate TNFR signaling capability. The ratio of p-IκB/IκB was calculated via densitometry.

**Supplementary Figure 3:**
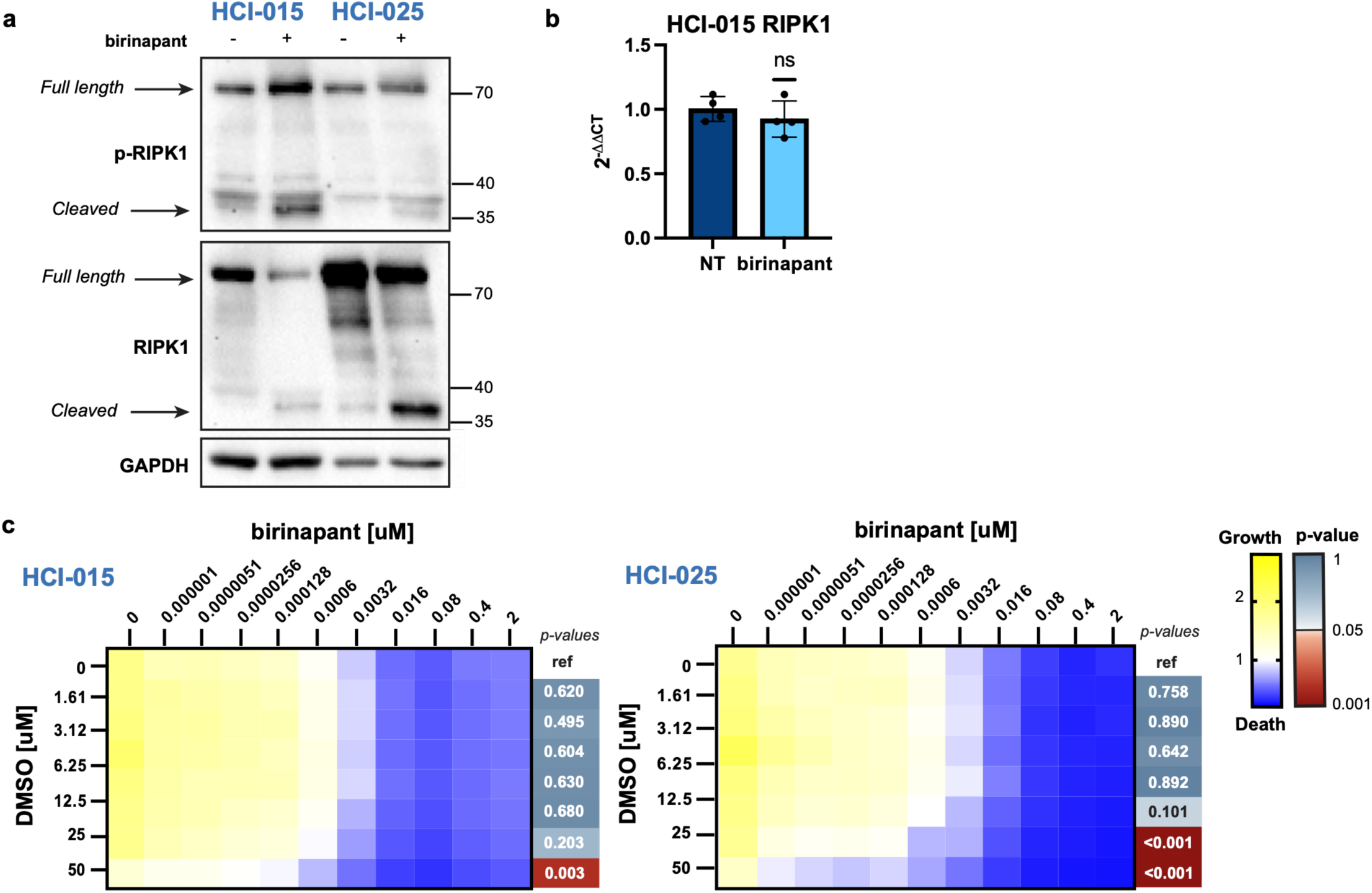
Birinapant induces RIPK1 protein cleavage but does not affect RIPK1 RNA levels in sensitive TNBC PDxOs. **(a)** S-PDxOs were treated with birinapant (10nM) in the presence of Z-VAD-FMK (25uM) for 12 hours. RIPK1 and p-RIPK1 were assessed via Western blot. The full length and cleaved fragment of RIPK1 are indicated. **(b)** RIPK1 RNA was quantified with qRT-PCR (n = 4 replicates) in the S-PDxO HCI-015 treated with birinapant (10nM) in the presence of Z-VAD-FMK (25uM) for 12 hours. Expression was normalized to GAPDH RNA and relative expression was calculated (2^-ΔΔCt^). Statistical significance was calculated using a two-sided paired Student’s t-test. Data are shown as mean +/- s.e.m. **(c)** S-PDxOs were pre-treated overnight with equivalent vehicle (DMSO) for Nec-1s used in **Fig. 3c** and subsequently treated with combination of vehicle and birinapant at the indicated concentrations for an additional 48 hours (3 biological replicates with 4 technical replicates each). For each biological replicate, the area under the curve (AUC) for each combination treatment row was compared with the corresponding AUC for birinapant alone (row 0; ref = reference) to determine the effects of DMSO on birinapant response. Statistical significance between AUCs was assessed using a two-sided paired Student’s t*-* test (n = 3 biological replicates). For all heatmaps, darker blue indicates cytotoxicity, white indicates a cytostatic response, and yellow indicates growth, and red p-values indicate statistical significance. While DMSO at high concentrations contributes to cell death in the presence of birinapant, Nec-1s did not.

**Supplementary Figure 4.**
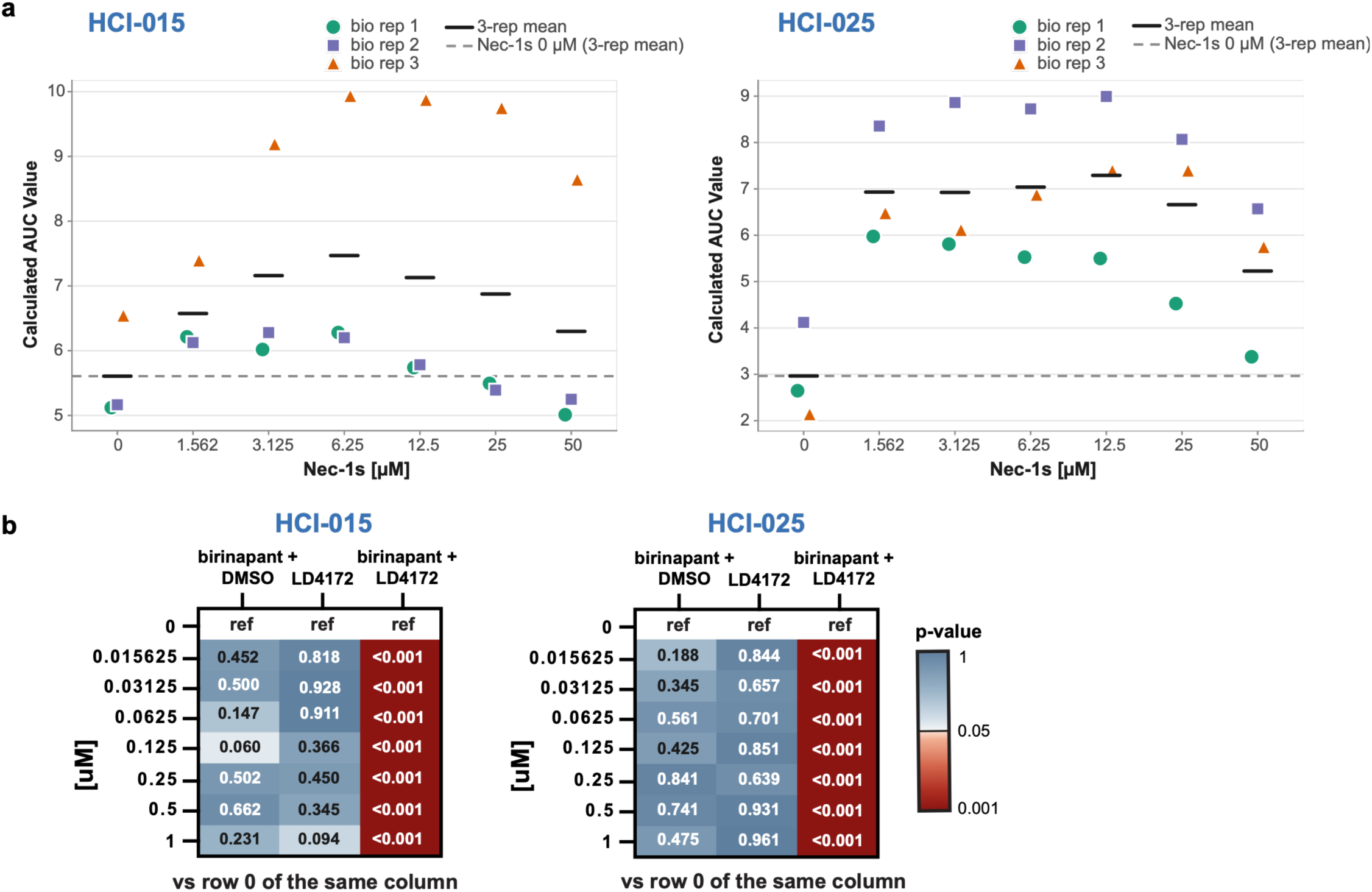
Depletion of RIPK1 protein protects against birinapant-mediated apoptosis. **(a)** Area under the curve (AUC) values calculated for each row corresponding to the drug-response heatmap in **Fig. 3c** to depict variance between biological replicates (bio rep, n = 3). The AUC replicate dot plots show the calculated AUCs at each concentration of Nec-1s treatment (μM; x-axis) to evaluate of the effects of Nec-1s addition on the birinapant dose-response curve. (solid black line = mean; dashed line = average AUC of row 0). **(b)** P-value heatmaps corresponding to the drug-response heatmaps shown in **Fig. 3e**. For each biological replicate, cell viability for each treatment concentration was compared to the corresponding non-treated control (row 0; ref = reference). Statistical significance between values was assessed using a two-sided paired Student’s *t*-test (n = 3 biological replicates), and p-values are color-coded by significance.

**Supplementary Figure 5:**
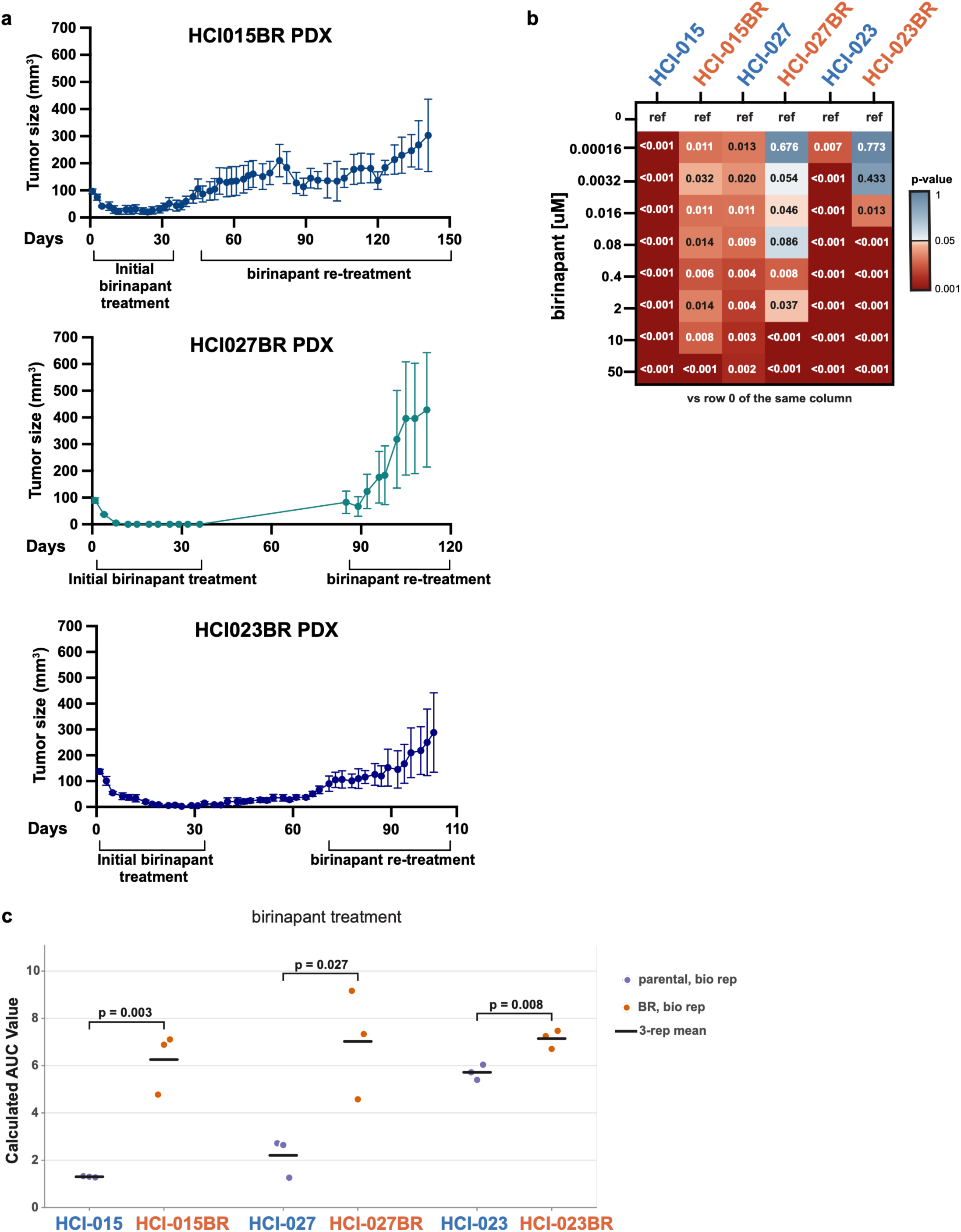
Development of PDxOs with acquired resistance to birinapant. **(a)** Mice with S-PDX (HCI-015, HCI-027, HCI-023) TNBC tumors were treated with birinapant (20 mg/kg, IP) three times a week for a 36-day period, starting when tumors were approximately 100 mm^3^. Tumors were then monitored off-treatment until they resumed growth, and re-treatment began when tumors regrew to 100 mm^3^. The birinapant-resistant (BR) tumors were harvested and established as new PDxO lines (HCI-015BR, HCI-027BR, and HCI-023BR). **(b)** P-value heat map corresponding to the drug-response heatmaps shown in **Fig. 4a**. For each biological replicate, cell viability for each treatment concentration was compared to the corresponding non-treated control (row 0; ref = reference). Statistical significance between values was assessed using a two-sided paired Student’s t-test (bio rep, n = 3), and p-values are color-coded by significance. **(c)** Area under the curve (AUC) values calculated for each biological replicate corresponding to the drug-response heatmap in **Fig. 4a** to depict variance between biological replicates and compare calculated AUCs. For each parental and matched BR PDxO, the AUCs of the three biological replicates were compared by an unpaired two-sided Student’s t-test, and p-values are indicated.

**Supplementary Figure 6:**
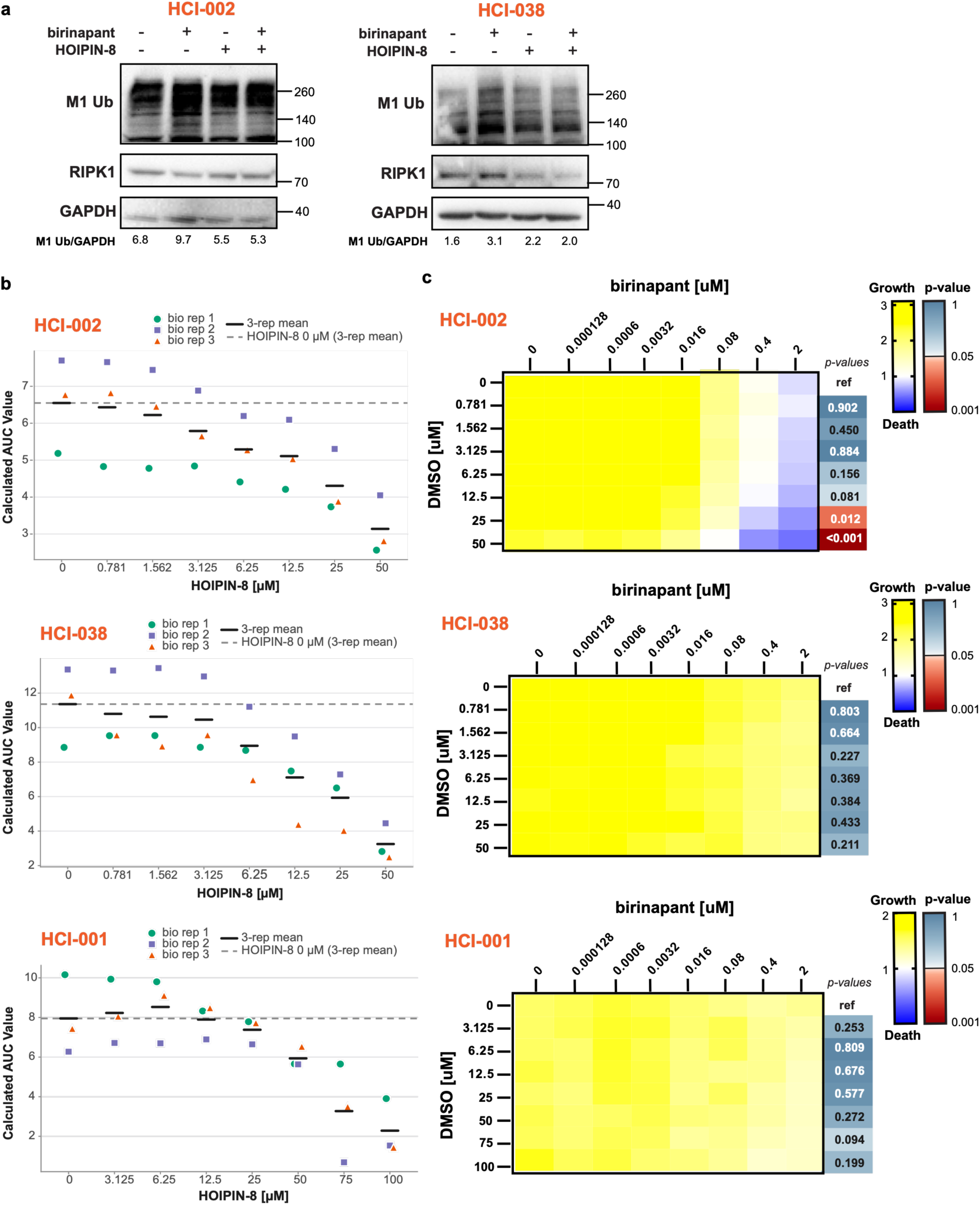
Birinapant treatment induces global M1 ubiquitination in resistant PDxOs. **(a)** TNBC R-PDxOs were treated with birinapant (10nM), HOIPIN (50uM), or the combination in the presence of Z-VAD-FMK (25uM) for 12 hours. Conditions that contained HOIPIN-8 were pre-treated for 12 hours (50uM). Abundance of M1 ubiquitin (M1 Ub) was assessed via Western blot. The ratio of M1 Ub to GAPDH was calculated by densitometry. **(b)** Area under the curve (AUC) values calculated for each row corresponding to the drug response heatmap in **Fig. 5d** to depict variance between biological replicates (bio rep, n = 3). The AUC replicate dot plots show the calculated AUCs at each concentration of HOIPIN-8 treatment (μM; x-axis) to evaluate the effects of HOIPIN-8 addition on the birinapant dose-response curve. (solid black line = mean; dashed line = average AUC of row 0). **(c)** TNBC R-PDxOs were pre-treated overnight with equivalent vehicle (DMSO) for HOIPIN-8 (shown in **Fig. 5**) and subsequently treated with combined vehicle and birinapant at the indicated concentrations for an additional 48 hours (3 biological replicates with 4 technical replicates each). Darker blue indicates cytotoxicity, white indicates a cytostatic response, and yellow indicates growth. For each biological replicate, the area under the curve (AUC) for each combination treatment row was compared with the corresponding AUC for birinapant alone (row 0; ref = reference) to determine the effects of DMSO on birinapant response. Statistical significance between AUCs was assessed using a two-sided paired Student’s t*-*test (n = 3 biological replicates), and p-values are color-coded by significance.

